# Experimental and computational analysis of REM sleep distributed cortical activity in mice

**DOI:** 10.1101/2023.01.19.524685

**Authors:** Mathias Peuvrier, Laura Fernandez, Sylvain Crochet, Alain Destexhe, Paul Salin

**Author notes:** (M. Peuvrier); (L. Fernandez); (S. Crochet); (A. Destexhe); (P. Salin). Equal contribution (Alain Destexhe, Paul Salin).

## Abstract

Although classically Rapid-Eye Movement (REM) sleep is thought to generate desynchronized activity similar to wakefulness, it was found that some brain regions can express Slow Wave activity (SWA), a pattern which is normally typical of slow-wave sleep. To investigate possible underlying mechanisms, we analyze experimental recordings and introduce a computational model of mice cerebral cortex in REM sleep. We characterized the patterns of slow-wave activity across somatosensory and motor areas, and find that the most prominent REM-related SWA is present in the primary (S1) and secondary (S2) somatosensory areas, more rarely seen in motor cortex, and absent from prefrontal cortex or hippocampus. The SWA also tends to be synchronized in S1 and S2. We next investigated possible mechanisms by using a computational model of the mouse brain consisting of adaptive Exponential (AdEx) mean-fields connected together according to the mouse connectome. To compare with experimental data, the local field potential is calculated in each mouse brain region. To reproduce the experiments, we had to assume a heterogeneous level of adaptation in different cortical regions during REM sleep. In these conditions, the model reproduces some of the experimental observations in the somato-motor areas and the other cortical areas. We then used the model to test how the presence of SWA affects cortical responsiveness. Indeed, we find that the areas expressing SWA have diminished evoked responses, which may participate to a diminished responsiveness during REM sleep.

## 1. Introduction

Classically, brain states such as wakefuless (W) and Rapid-Eye Movement (REM) sleep are thought to generate essentially desynchronized activity, while slow-wave sleep (SWS) typically generates synchronized slow oscillatory waves (Niedermeyer, 2004). However, it was found recently that slow-wave activity (SWA) can also be seen during quiet wakefulness in some brain areas (Crochet and Petersen, 2006; Vyazovskiy et al., 2011; Zagha et al., 2013; Fernandez et al., 2016). SWA can also be observed in REM sleep, in human (Baird et al., 2018; Bernardi et al., 2019), and in mice(Funk et al., 2016; Timofeev and Chauvette, 2019). In their study, Funk et al showed local SWA in the superficial layers of primary areas of the mouse sensory cortex, while the activity in secondary sensory and associative cortical areas seemed desynchronized (Funk et al., 2016). These results may suggest that SWA would be present in all primary sensory areas and filter sensory information during REM sleep.

### Experimental recording of slow-wave activity

The slow waves of SWS are composed of delta oscillations of 2-4 Hz and the slow oscillations (SO <2 Hz). This slow oscillatory activity corresponds to the coordinated spiking activity of a large population of neurons, alternating between active (upstates) and silent (downstates) periods. Taken together, these oscillations make up the slow wave activity (SWA) which reflect the homeostatic processes of sleep. Thus with an increase of the sleep pressure, the power of SWA increases and conversely when the sleep pressure decreases the SWA power decreases. Several studies suggest that the slow and delta oscillations observed are generated in the neocortex (Sanchez-Vives and McCormick, 2000; Timofeev et al., 2000). Brain activity during SWS is accompanied by a phasic activity, spindles, which are brief and of rapid frequency (1s, 10-14Hz) (Steriade and Timofeev, 2003).

In contrast, REM activity is associated with low amplitude higher frequency theta (5-10 Hz) and gamma (35-80Hz) activity (Mizuseki et al., 2012). Thus the brain states of SWS and REM examined with EEG recordings appear entirely opposite. It is only recently that phasic events of faster theta with strong gamma coupling were observed in the hippocampus (Montgomery et al., 2008; Brankačk et al., 2012; Scheffer-Teixeira et al., 2012). Previous work on cat associated increase in theta frequency and power with PGO, rapid eye movement and other body phasic activities (Karashima et al., 2005; Lerma and Garcia-Austt, 1985; Robinson et al., 1977; Rowe et al., 1999; Sei and Morita, 1996; Sanchez-Vives, 2020). Montgomery et al hypothesise that phasic bouts may provide windows of opportunity for the transfer of information between hippocampal areas, opening new ways to consider phasic cortical events with specific functional implications in REM.

Advances in in vivo electrophysiological techniques (intracellular recordings, multisite local field potential (LFP) and spike recordings and high-density EEG recording) have revealed that in the neocortex W is characterised by substates alternating SWA and desynchronized activity during quiet and active wake respectively. These SWA in quiet wake share some common features with delta oscillations in SWS and are generated in sensory, motor and associative cortical areas (Crochet and Petersen, 2006; Vyazovskiy et al., 2011; Zagha et al., 2013; Fernandez et al., 2016). Intriguingly, SWA is also observed in REM in LFP recordings in the superficial layers of primary areas of the mouse sensory cortex (Funk et al., 2016). In contrast, the activity of the secondary sensory, motor and associative cortical areas would be desynchronized as in EEG and hippocampal recordings (Funk et al., 2016; Montgomery et al., 2008). This finding seems to be opposed to EEG work showing that REM activity is characterised by rapid oscillations of low amplitude. However, the results of Funk’s study may help explain the REM “paradox”: an EEG activity close to wakefulness that is associated with high sensory threshold due to synchronised slow activities in the primary sensory cortices. If SWA are present in these primary cortical areas they could filter sensory perception during REM like during SWS. Thus, this finding leads to reconsider the importance of SWA in REM as compared to SWS and W.

Very few studies have characterised the differences between SWA in REM and those in SWS and W. A detailed analysis of cortical SWA in REM versus SWS and wakefulness has not yet been performed. Are REM slow waves different in amplitude and shape compared to SWS and W? What makes this slow activity possible? Is it modulated by the phasic theta events previously identified? By recording LFP activity in several cortical areas using multi site recordings it becomes possible to directly address these questions and to determine similarities and differences in the scale of the individual slow wave as well as in the temporal and spatial distribution of SWA in REM compared to SWS and W.

Using multisite LFP recordings, we first analyse REM SWA in comparison to the SWA of SWS and W in the mouse somato-motor cortex. Since this SWA is important in this cortex, we examined whether it is correlated between these cortical areas on the one hand, but we also determined whether it is correlated to other areas where faster oscillations are predominant such as the hippocampus. We show that REM SWA is characterised by Delta waves in primary and secondary cortical somatosensory (S1 and S2 respectively), as well as in primary motor areas (M1), and that SWA alternates with faster low voltage oscillations during REM episodes. This indicates a coordination in the different cortical regimes in place, and gives us new insights into the brain capacity for information integration in REM.

### Computational models of sleep

Until now, most modeling efforts regarding REM aimed at building models concerning the genesis, maintenance or regulation of this sleep state in relation to the other states of vigilance (McCarley, 2004; Héricé et al., 2019). Surprisingly, little has been done to model cortical activity during REM, compared to the numerous computational works concerning SWS. Models of cortical dynamics of sleep (and wakefulness) with fine temporal precision rely on neuronal network models with single neurons, of various degrees of detail. Those models, and the thalamocortical models in particular, were very fruitful in describing the slow oscillatory Up and Down regime of the SWS (Bazhenov et al., 2002; Destexhe et al., 1999; Compte et al., 2003; Hill and Tononi, 2005). In such models the neocortex is composed of a few thousand neurons that would represent one to two cortical columns, with a specific connectivity, often inspired by biological observations (Markram et al., 2004; Lübke and Feldmeyer, 2007). However, expanding such models to the whole cortex would require a lot (way too much) of computational power, making it almost impossible to run, but also extremely difficult to analyse. This is why we looked at population models, abstracting the neuron activity to drastically reduce the computational cost, while remaining completely relevant and very detailed in regard to experimental LFP recordings. It is a mesoscopic approach that describes the average collective properties of neuronal populations, instead of simulating each individual neuronal dynamics(Deco et al., 2008).

Furthermore, this modeling approach is inspired by the work of Goldman et al. (2020) that produced important features of global cortical dynamics comparable to W and SWS in human brain-scale simulation. Using The Virtual Brain (TVB) simulator (https://www.thevirtualbrain.org/tvb) to integrate population models as the nodes of a network in which the connections are based on brain connectivity as defined from diffusion imaging experiments. In our model we will use the TVB mouse counterpart, The Virtual Mouse Brain (TVMB; Melozzi et al. (2017)). TVMB can construct individual mouse brains from dMRI experiments or from tracer experiments realized by the Allen Institute. Thanks to that we could use a detailed connectome of the mouse cortex that fit well the level of description we have in our biological dataset.

To model neuronal populations, we used an AdEx meanfield model, as described in di Volo et al. (2019). This biologically realistic mesoscale model was built to correctly predict the level of spontaneous activity observed in AdEx spiking neural networks with Asynchronous Irregular (AI) or Up and Down regimes. Two network regimes that are somehow the computational analogs of biological Wakefulness and SWS sleep respectively (Destexhe, 2009). In those models, the spike-triggered adaptation parameter is sufficient to switch from one regime to the other. It is interesting to note that this adaptation parameter is strongly related to the type of adaptation produced by high-threshold calcium-gated current (Augustin et al., 2013). This current is known to mediate the spike frequency adaptation in cortical pyramidal neurons but also to be deactivated by acetylcholine (Mccormick, 1992). Therefore it is not surprising that in models with strong spike-triggered adaptation, the adaptation mediated by this current is strong, such as in a cortex without acetylcholine signaling, such as in SWS.

Using this model of the whole mouse cortex, we can simulate the transition from a Wake model (W-model) to a SWS model (SWS-model) by changing only the spike-triggered adaptation parameter. Yet there is no REM-model. Here we present a whole-brain model that we tuned to fit LFP recording of mouse REM sleep. In particular, we will focus on cortical dynamics in the somatosensory primary and secondary cortices (S1 and S2), the primary motor cortex (M1), the medial prefrontal cortex (mPFC) and the hippocampus (dCA1), with a particular interest regarding the REM SWA as recently observed (Funk et al., 2016; Bernardi et al., 2019; Soltani et al., 2019). With these considerations to build our REM-model, we propose a simple topological organisation of the spike-triggered adaptation parameter in the model. This organisation allows the simulation of different patterns of activity across the cortex that reproduce a part of the diversity of activities we observed experimentally. Finally we also use this new model to predict the potential cortical response to cortical stimulation in this state.

## 2. Methods

### 2.1. Electrophysiological signal analysis

The data analyzed in the present study come from a database acquired in unanesthetized head restrained mice recorded by L.F. and S C. Part of this dataset has been recently analyzed and published to characterize the cortical slow waves of wakefulness and SWS (Fernandez et al., 2016).

#### Animal preparation and surgery

Briefly, nineteen C57BL/6 male mice (Janvier SAS, St. Berthevin, France), aged of 6 to 8 weeks at the time of surgery, were used for the study. All procedures were approved by the University of Lyon 1 Animal Care Committee (project DR2013-4) and were conducted in accordance with the French and European Community guidelines for the use of research animals.

The data of only seven out of the nineteen mice was analyzed here. LFP for those mice were recorded at the following sites: the barrel field of the primary Somatosensory cortex (S1), the secondary Somatosensory cortex (S2), the primary Motor area (M1), the prelimbic area of the medial prefrontal cortex (mPFC), and the area CA1 of the dorsal hippocampus (dCA1).

In addition to the LFP microelectrodes, two surface EEG were implanted over the parietal (AP 2.0, lat 1.5) and frontal areas (AP 5.3, lat 1.5) onto the dura mater of the contralateral hemisphere, two EMG electrodes were inserted bilaterally in the neck muscles, and two silver wires were inserted on both sides of the cerebellum for reference and grounding. Then, to allow for painless head-fixation during recording sessions, a light-weight metal head-post was cemented to the skull.

**Table 1.** LFP implantation coordinates. In bold is highlighted the areas that will be studied in this article.

| Area | S1 | S2 | M1 | A1 | V1 | PtA | mPFC | dCA1 |
| --- | --- | --- | --- | --- | --- | --- | --- | --- |
| Antero-Posterior | 2.8 | 3.1 | 5.1 | 1.3 | 0.5 | 1.8 | 5.7 | 1.3 |
| Lateral | 3.2 | 4.0 | 1.8 | 4.0 | 2.5 | 1.8 | 0.3 | 2.0 |
| Depth | 0.5 | 0.8 | 0.4 | 0.6 | 0.4 | 0.4 | 1.85 | 1.3 |

#### Habituation and recording

Habituation and recording sessions were performed between 12AM and 5PM, during the light period in a semidark light condition, with ambient room temperature (22±2°C), in an electrically shielded recording chamber.

After recovery from surgery, mice were progressively habituated to head-restraint conditions with repetitive daily training. The recording sessions started when the first SWS episodes were observed.

#### Sleep scoring

Epochs of wakefulness, SWS and REM were scored using EEG and EMG. REM sleep was identified by theta oscillations on the EEG and neck muscle atonia on the EMG. Artifacts, drowsiness periods and intermediate sleep were discarded. Therefore, each episode of vigilance states (W, SWS, REM) showed a clear separation.

#### LFP dataset

For each mouse we have sets of brain state episodes, determined with the sleep scoring, corresponding to specific brain states with the following data: EEG, EMG and a set of 5 to 6 LFP (all sampled at 1kHz). LFPs implanted in each mouse corresponded to the configuration presented in the previous table. In this study, we only use the first configuration, with 7 mouse recordings in: S1, S2, M1, mPFC, dCA1. This allows the study of interactions between S1 and other areas of the somatomotor cortex. Quantitatively this database includes 341 episodes of W (total duration of 14866 s, mean episode duration: 43.6 ± 36.05), 318 episodes of SWS (total duration 12274.08s; mean: 38.6 ± 25.1s) and 120 episodes of REM (total duration 7418.68s (mean: 61.82 ± 40.1s).

#### Spectral analysis

To compute the spectrogram of LFP, from the dataset or from the model, we used the Welch method. The power spectral density is estimated using a Hamming window of 4 seconds with 2 seconds overlapping, with the exception of 1000ms windows with 300ms overlap for the phasic theta analysis. The power spectrum is expressed in V^2^. One spectrogram is computed for each brain state episode recorded, and they are used to compute band power and to find frequency peaks in the delta band.

The maximum frequency peak is the highest peak in the frequency range. The absolute delta power is the integral of the area, between the frequency bands and below the periodogram line. To integrate this area, we approximate it using the composite Simpson’s rule. The idea behind it is that the area is decomposed into several parabola and then summed up their area.

#### Delta activity in the signal

In order to understand the delta activity, we first detect the individual delta cycles in the signals. Then we looked for delta bouts, defined as short periods (from 1 to 10 seconds) with important density of delta cycles. The different steps of this analysis are shown fig S1, where we have the raw signal, the delta cycle identification and the delta bout determination.

#### Bicycle algorithm

To detect and analyze Delta cycles in our data, we used a method inspired and based on the Bicycle algorithm (for cycle-by-cycle analysis) and python library (Cole and Voytek, 2018). In this method, the LFP signal is segmented in cycle periods in a few. The signal is bandpass filtered, and the zero-crossings are indexed, then, on the raw data peaks and troughs were found as the minima and maxima between two zero-crossing time points. Finally, flank midpoints are the point halfway between the peak and trough voltage. Extremas and flank midpoints allows us to compute various cycle features such as the period, the peak and trough duration or voltage, the rise-decay, and peak-trough symmetries. The instantaneous phase is estimated by interpolating between extrema (the peaks and troughs) and flank midpoints. In the original use of the bicycle algorithm, it is proposed to validate a putative oscillatory cycle when it is surrounded by similar cycles, taking part into a larger, sustained oscillatory bout. However, delta cycles were frequently interspersed with faster activities, and we observe isolated delta cycles. Thus, we used permissive settings from the original library, but added another threshold as compensation and redefined the detection of Delta bouts.

#### Determination of delta cycle

To find delta cycles we used the algorithm to compute cycles from trough to trough, centered on the peak with a narrow bandpass filter between [2 and 4.5 Hz] for the first step. Then, using a simple amplitude threshold, cycles with amplitude high enough, compared to the rest of the episode signal, were classified as putative delta (fig sup S1, blue line) without regard to other detected cycles. This threshold was not restrictive at all and captured many false positives. But it was only used to calculate the actual amplitude threshold, as the median amplitude value for all putative delta cycles found in S1 and S2 during SWS for the specified individual This second decision parameter (fig sup S1, purple line) restrict the delta cycles to those above 80% of the amplitude of the Slow wave sleep Delta cycles recorded in the primary and secondary somatosensory areas for each mouse. The cycle-by-cycle analysis of delta oscillations has been progressively optimized and verified by direct observation of the dataset by 2 authors (MP, PS). While this method was effective at detecting a significant number of Delta cycles in the signal, it still missed few (usually isolated) delta cycles.

#### Delta bouts

To understand the repartition of delta cycle in the signal, we categorized the LFP signals in delta and non-delta bouts. Those bouts are short periods (usually about 2 to 4 seconds) of signal characterized by important Delta activity, or its absence. To determine the temporal distribution of Delta, we describe LFP signals in delta and non-delta bouts. Validated delta cycles that are close enough were gathered together, adding non detected cycles if they were surrounded by delta cycles and that at least 75% of the bout is made of validated delta cycles. Then those periods were validated as delta bouts if they lasted at least 1 second and were constituted of at least 3 delta cycles (fig sup S1, bold green). Non-delta bouts are periods of at least 1 second without any validated delta cycle. Thus, isolated delta cycles and very short non-delta portion of signal are excluded of this categorization as bouts.

### 2.2. Whole brain modeling

The whole brain models we discuss are built at the neuronal population resolution and implemented in The Virtual Mouse Brain. It consists of a network of AdEx mean fields connected together following Connectome-based information from the brain Allen institute (https://alleninstitute.org/). In order to keep this section as short and simple as possible, we’ll spare some mathematical details regarding the mean-field model or the TVB implementation. The corresponding informations could be found in di Volo et al. (2019) and Goldman et al. (2020, 2022) respectively.

All simulations presented in the paper were run for ten seconds, with the first two seconds (the simulation initiation) being systematically excluded. The results presented for the brain state model in the figure 4 are results from a single simulation, while results concerning the stimulation part (Fig.5) are the average of 10 simulations (See the related part of the method).

#### The Virtual Mouse Brain

We use the simulation framework The Virtual Mouse Brain (https://www.thevirtualbrain.org/tvb) to perform detailed connectome-based mouse brain simulations. With the Allen Connectivity Builder pipeline, experimental connectivity data from dMRI or tracer experiments are used to build a precise mouse connectome. The connectome is a square matrix (origin to target structure and vice versa) informed with connection strength and tract length. The Allen Connectivity Builder also provides a 3D matrix which represents the volume of the mouse brain to create a region volume mapping (Melozzi et al., 2017). This framework allows us to virtualize a mouse brain with a population model at the nodes corresponding to a cortical column or a brain area, and edges corresponding to connection strength between regions. We use a connectome composed of 104 cortical nodes Rabuffo et al. (2021). We selected this connectome over more detailed ones that includes thalamus and other subcortical nuclei for two principal reasons. First we don’t have simultaneous recordings including subcortical areas in our dataset. Second, the population model we decided to use was built to match the dynamics of cortical spiking neuronal network models. Therefore more complex connectomic dataset are not more relevant regarding our biological dataset while it could induce some errors in the network dynamics.

#### Mean field model

The population model used is an AdEx mean-field model as described by di Volo et al. (2019). This mean-field model, based on a previous Master Equation formalism (El Boustani and Destexhe, 2009) was built to match the spontaneous activity of networks of 10000 Regular and Fast Spiking AdEx (Adaptive Exponential) neurons. It takes into account the nonlinear effects of the conductance-based synaptic interactions as well as the spike-frequency adaptation of excitatory regular spiking neurons. The mean-field has two neuron populations with different intrinsic properties. One is excitatory, called the “regular spiking” (RS), and is affected by the spike-frequency adaptation, as cortical pyramidal neurons of the somatomotor cortex. The other is an inhibitory population called “fast spiking” due to its usually higher firing frequency as cortical parvalbumin GABAergic interneurons. The dynamics of the two population firing rate and their covariance are computed as a function of the transfer function (TF; the firing rate of one population as function of the inputs it receives), which is itself dependent on the population adaptation variable W. This population adaptation has two important parameters: the sub-threshold adaptation *a* and the spike-triggered adaptation *b*. It should be noted that this parameter *b* affects only the excitatory population. Another difference in the dynamics of the two populations is the inputs considered in their transfer function. Indeed, only the excitatory population is affected by external noisy drive, while both populations are affected by external inputs (such as thalamic inputs).

This mean field describes the population dynamics of a single, isolated cortical column. The parameters used in our study are the same as in previous ones. They produce biologically realistic and sustained brain dynamics. With those parameters we can switch from an AI to an Up and Down model regime changing only the spike triggered adaptation parameter *b*. Therefore, except this single parameter *b*, all parameters are the same in every node and every simulation presented.

#### Networks of mean-field models

In order to model large brain regions, we represent interconnected cortical columns as a network of mean-field models. The inter-cortical mean-fields connections are done by excitatory interactions only, while inhibitory connections remain locally within the column.

Therefore, once in the network, the dynamics of the single mean-field is changed as it is now impacted by the excitatory input exerted from multiple excitatory populations. The integration of all excitatory inputs is computed using the distance between the origin to target node and the axonal propagation speed to account for the delay of axonal propagation. Thus it is at this step that the connectomics information impacts and shapes the whole model dynamics.

#### REM model spike-triggered adaptation parameter

The REM model tries to account for the diversity of activity observed experimentally. We look to produce a simulation in which some areas marked by a Delta activity intertwined with periods of faster oscillations (in different proportions according to the area), while other areas would be closer to an ‘Awake-like’ activity. We hypothesize that regional variation of adaptation could be a key to get such a model as it is the parameter used to switch from Wake like to SWS like in the reference model.

In order to keep the model easy to use and describe, we present a model with only three different values of adaptation. This number is the smallest number of different values we found that produces a satisfying result. The different adaptation values used have a conductance of 5 nS (as for our reference W-model), 30nS and 60nS (as for our reference SWS-model). We decide to spatially arrange the adaptation to match with the observation of the decreasing prevalence of delta activity in the LFP going from S1 to S2 and to M1. We place the maximum adaptation level (60nS) centered at S1 barrel cortex.At 12mm from this center (encompassing S2) the adaptation was decreased to 30nS, and then, 12mm further away (encompassing M1) it is decreased to 5nS. We add some exceptions to this method of application of the b parameter regarding hippocampal (field CA and dentate gyrus) and prefrontal (anterior cingulate and prelimbic areas) areas that were parameterized with b = 5nS disregarding their position in the model. We think that this setup is more in accordance with the literature (Mizuseki et al., 2012; Maciel et al., 2021), although we did not look further into optimizing the activity of those regions.

#### REM shuffled as control

With our REM model we introduce a mirror model that was used as a control: the REM shuffled. This model is built to have exactly the same number of nodes with the different values of adaptation. However in this model the topological arrangement is randomized, losing the spatial features of the model for the adaptation. The goal of this model is to look at the importance of this spatial arrangement of adaptation parameter *b*. We use it to question if the phenomenon need the organized recruitment of multiple nodes, or if any node with high enough adaptation could produce the delta activity disregarding (or with randomized) the average input of the other cortical areas. Most results concerning this model are not shown to keep the results clear, although it should be considered that the results of those REM shuffled models are always more variable than any other model.

#### LFP measurement

The estimate of the LFP signal of each node is computed with a Kernel-based method based on previous work (Telenczuk et al., 2020; Tesler et al., 2022). First demonstrated in a neuron network and then extended to mean-field models, it relates the spiking of neurons to the LFP through efferent synaptic connections. The LFP was calculated as the convolution of the unitary LFP waveforms modeled from the spiking activity of the field. This method also takes into account the relative contribution of the neuronal population to the signal according to the distance between the origin of the uLFP and the recording site.

In this paper the LFP signal was simulated as if originating from an electrode placed in a superficial layer of the cortex thus corresponding to the location of the electrodes in the database. Thus the signal is inverted compared to the population firing rate. The LFP is calculated from the firing rate of each node independently, without horizontal contribution from other nodes which could have accounted for some volume conduction effects

#### Stimulation

With TVMB we used an external input in order to stimulate our model. The stimuli we used consist of a single pulse of 20ms produced by an external and repeated every 1.25sec (±250ms) and starting 3 seconds after the beginning of the simulation Fig. S4).

This external node is connected to it’s target with a specific connection weight that allows us to control the strength of the stimulus. Different connection weights were tested by stimulating S1 in our different state models. Looking at the kinetics of the evoked potential we found a point where the time to the first peak response is small and remains stable in all conditions for connection strength of 10-3 and above. Therefore all results presented in the main article have the same connection strength of 10-3.

The stimulation experiment consists of a set of ten simulations in which the stimuli is provided to one node only (ie simulating the electrical stimulation of neurons in a cortical area), from the right hemisphere, with 5 or 6 stimulus repetition. For each we therefore have a set of above 50 Stimulus Evoked Potentials (SEP). We performed this stimulation experiments for the 5 areas we focus on this article (S1, S2, M1, mPFC (its equivalent is the anterior cingulate in the model), dCA1) in each model state (AI, UpDown, REM-model, REM-shuffled). The results presented is the average of the SEP obtained in each condition.

## 3. Results

We begin by analyzing the occurrence and distribution of slow-waves in REM sleep and then introduce a computational model to account for these results.

### 3.1. Analysis of slow-waves in REM sleep

#### Primary somatosensory area signal in different brain states

Figure 1 shows the three states of the wake-sleep cycle in mice, wakefulness (W), slow-wave sleep (SWS) and rapid eye movement (REM) sleep recorded by local field potentials (LFP) in the mouse primary somatosensory area (S1). W and SWS classically display two drastically different activity states, asynchronous and synchronized slow-waves respectively as in EEG recordings (data not shown). During REM sleep, however, one can see that the situation is less clear. While asynchronous activity usually dominates in EEG and in some LFP recordings of cortical areas, other brain areas exhibit local slow-wave activity, in middle and superficial layers of primary cortices, as pointed out previously (Funk et al., 2016). The differences between these states is also apparent in the power spectral density (PSD) of the LFP signals (Fig. 1B). Here, the broad low-frequency peak, culminating in the Slow Oscillation (SO) band (<1.5Hz), during SWS contrasts with the absence of clear peak in W, while in REM, a low-frequency peak appears around 2-4 Hz (Fig. 1C.). The REM slow-waves seem thus faster than the classic slow wave complexes of SWS. This difference in frequency is characterized further in respect to the Delta component of the slow activity. Traditionally considered characteristic of SWS, the power in the delta frequency band was very similar, if not higher, in REM [Fig. 1D.). However, the shape of the Delta cycles detected on S1 LFP signals was different in REM compared to SWS (Fig. 1E.). In regard to the average rising and decay duration of the Delta cycles, the delta rise time was longer for REM oscillations than for SWS oscillations (N delta cycles in S1 during SWS = 15963 cycles; REM = 11171 cycles. Delta mean rise time: REM = 185.43±13.06ms, SWS = 167.66±12.35ms student t value = 13.17, p ≈ 1e-33), while delta decay was shorter for REM oscillation than for SWS oscillations (Delta mean decay time: REM = 131.09± 11.9ms, SWS = 146.19±11.80ms student t value = -11.89, p ≈ 2e-28). Therefore there was a clear difference in the ratio between rise and decay time of Delta cycles in REM compared to SWS. As illustrated in Fig. 1F the distribution of rise and decay time of Delta cycles in the two sleep states overlap poorly, and REM delta was more biased toward high rising time and low decay time. Rise and decay time of delta cycles that were occasionally observed during W (i.e. quiet wake) were shorter than those of the SWS and REM. It should be noted that our algorithm was not tuned to optimize the detection of W delta, and thus might be less relevant than what we find for REM sleep.

**Figure 1:**
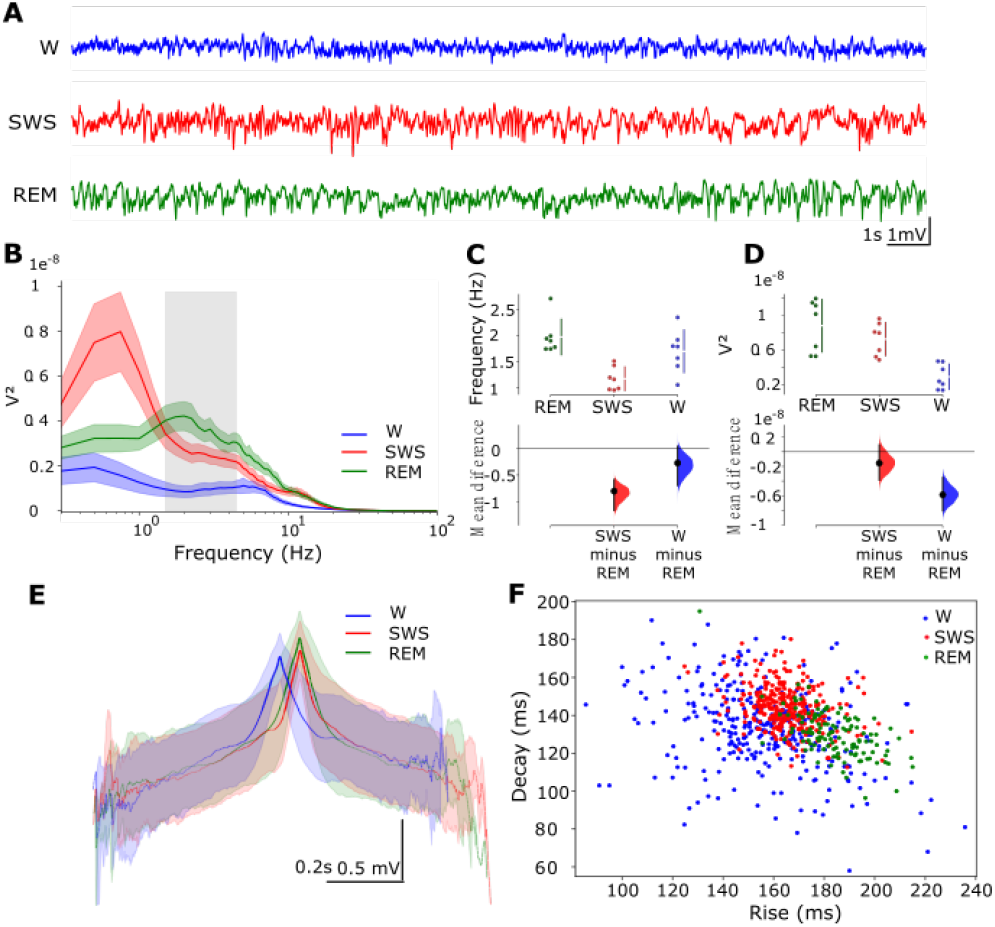
Delta in the somatosensory cortex across brain states. (figure A to G colour code: blue for W, red for SWS and green for REM). (A) Twenty seconds of LFP examples recorded in area S1 during sleep and wakefulness. (B) Mean power spectrum of S1 recordings. The grey area corresponds to the delta band, between 1.5 and 4.5 Hz. (C) Average frequency of the highest peak in SWA band (0.5,4.5Hz) Each point corresponds to the mean, for one subject of the frequency peak found in the power spectrum. (D) Delta band (1.5-4.5 Hz) power amplitude in S1 across states. delta bandpower is shown for each subject, with colours corresponding to the brain state. (E) Average Delta cycle detected in S1 for each state. The ratio Ndatapoint/totalNcycles is reported as the alpha(intensity) of the line plotted (with a minimal value when less than 30% of the cycles are involved). (F) Scatter plot of Delta cycles rise and decay time in S1 across states: Each dot corresponds to the mean rise and decay time of delta cycles detected during one episode of the three brain states This figure shows the dispersion of Delta cycles rise-decay symmetry.

#### Delta waves in different cortical areas in REM

To investigate the spatial distribution of REM slow-waves, we next compared LFP recordings in the different cortical areas during REM sleep.

Figure 2 shows the typical LFP activity observed during REM in simultaneously recorded cortical areas. Important differences in the amplitude as well as in the oscillation rhythms were observed. The area S1 and S2 presented bouts of large amplitude delta oscillations, intermingled with periods of smaller and faster signal fluctuations. In the area M1 we also found some large amplitude Delta waves, but sparser than those of the somatosensory areas, and most of the signal is of small amplitude with no clear prevalent oscillation. Finally, the LFP of the dCA1 area of the hippocampus showed a sustained theta oscillation along the REM episodes, which is a typical signature of REM in rodents (Buzsáki et al., 2003). Almost no delta oscillations were found in the dCA1 LFP. The spectral analysis of LFP confirmed these differences between the activities of the cortical areas recorded (Fig 2.B). The spectra of S1 and S2 were very similar with a clear peak in the Delta frequencies. In the area M1 there was also a delta peak above the 1/f trend of the LFP spectrum but its amplitude was considerably smaller than that of the somatosensory areas. LFP spectral analysis of the area mPFC did not have a delta peak and and therefore the analysis of delta oscillations in this area was not pursued. On panels D and E of the figure 2 were shown the averaged Delta cycles centered by their peak, and the distribution of the rise and decay duration of those Delta cycles. From those, it appears that the delta cycles found in S1 and S2 were extremely similar, with a bias toward long rising and short decay time, while Delta cycles from M1 were less stereotyped. They had a tendency to have a longer time decay, which might be due to the fact that Delta events in M1 were much more isolated and rarely immediately followed by another event.

**Table 2.** Percentage of somatomotor REM sleep LFP signal characterized by Delta bouts and Non-Delta bouts.

|  | S1 | S2 | M1 |
| --- | --- | --- | --- |
| Delta bouts | 47.33% | 30.373% | 7.362% |
| Non-Delta bouts | 41.441% | 53.25% | 84.283% |
| Mean Delta bout duration | 2.951s | 2.494s | 2.006s |

**Figure 2:**
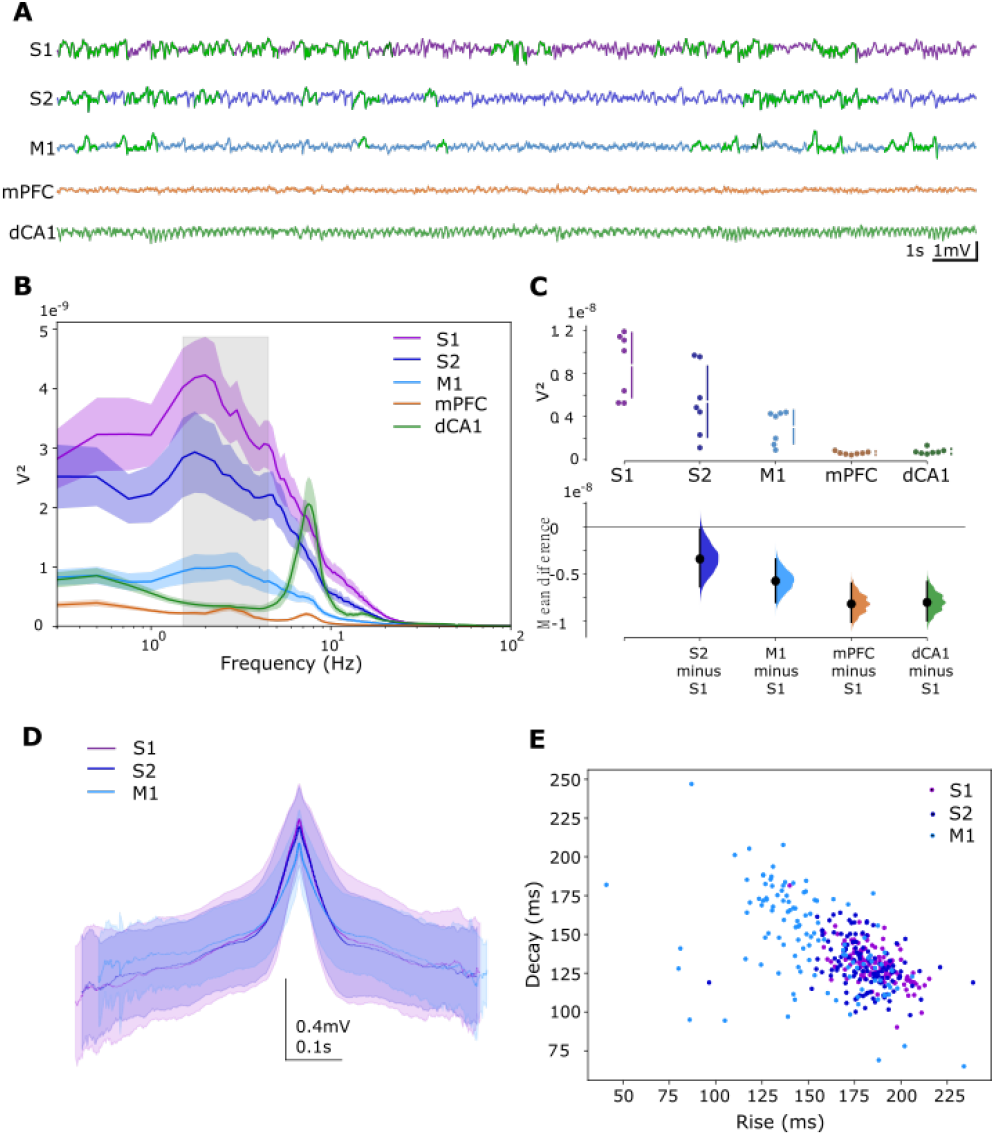
REM sleep delta oscillation in the cortex. (Figure colours: Purple: S1, dark blue: S2, light blue: M1, green: dCA1, orange: mPFC) A) 20 seconds of simultaneous LFP recording in 5 different areas during REM. Signals are shown with a common scale. Delta cycles detected in the signal are highlighted in green. (B) Average Power spectrum for the different areas for all REM episodes. Grey square shows the 1.5-4.5Hz delta band. (C) Delta band power, averaged for each mouse, in the different areas. Below is the Delta power compared to its distribution in S1. (D) Representation of the mean delta cycles, centred around their peak, over all detected cycles in the somatosensory and motor area. Line intensities represent the portion of delta cycle from the total pull recorded at specific area used for each datapoint. (Approximate number of cycles considered in each area: S1: 10000, S2: 8000, M1: 5000). (E) Scatterplot of the rise, decay symmetry of delta cycles in the somatosensory and motor area. Each dot corresponds to the mean values for delta cycles recorded in a REM episode.

#### Temporal distribution of REM-sleep delta bouts

Next, to determine the temporal distribution of REM-sleep delta waves, we detected successive delta cycles and found that periods of delta waves alternate with periods of asynchronous activity which we characterized in Fig. 3 as the delta and non-delta bouts.

**Figure 3:**
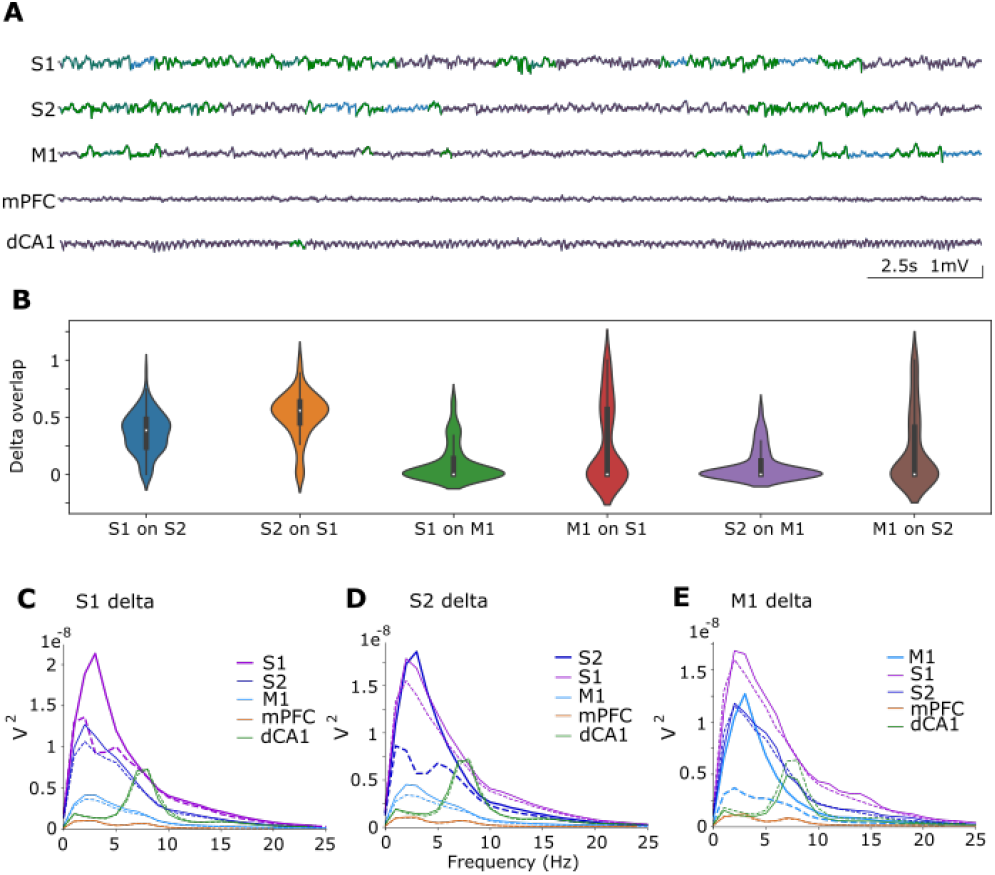
Temporal presence of Delta in REM. A) Same LFP recording as shown before. Colours corresponding to the delta chunk analysis result. Isolated delta cycles are in light green, Delta bouts in bold green and non-Delta bouts in red. In blue are the non-scored & non-delta parts. (B) Delta bouts overlap between areas. On the y axis is the percentage overlap of delta bouts from the origin to the target (S1 on S2: % of S1 delta bouts with simultaneous Delta bouts in S2). Values are scaled between 0 and 1 (100% overlap). For each directed pair of areas the distribution of this overlap measure calculated for every delta chunk period of the origin area is shown in the form of a violinplot to illustrate data variability. (C) we analysed only the areas of the somato-motor cortex. Power spectrum of S1 Delta bouts and Non-Delta bouts. For each area (with the same colour code as in figure2), the power spectrum shown is the average power spectrum of this area, for the Delta chunk periods of S1 in plain line, and for the non-Delta periods of S1 in dashed line. The thickest line highlights S1, the area from where the Delta / non-Delta periods are determined for this figure. (D) Same as C centred on S2. (E) Same as C centred on M1.

Our detection of Delta bouts allows us to discriminate periods (of at least a second) that are dominated by Delta activity from periods without any Delta activity, while excluding periods that would be considered as mixed-activity. With this approach we can describe at least 80% of any of our REM signals. This can be seen in Figure 3.A as the thick green and red lines, that represent Delta bouts and Non-Delta bouts respectively, occupy the major part of the signals. While in S1 the Delta bouts occupy almost half of REM episodes and last about 2.9seconds, it describes only 7.36% of the LFP signal of M1 in REM and lasts only 2 seconds on average. The result is intermediate for S2, where Delta bouts describe a large part (about 30%) of the signal and mean duration of the Delta bouts is 2.49sec in this area.

Next, we determined if Delta bouts occurred simultaneously in the different cortical areas. We thus assessed the level of temporal overlap of the delta bouts in the area S1, S2 and M1. For reference we look at the overlap of S1 on S2, the distribution peak was around 0.5, meaning that for about 50% of S1 Delta bouts duration, we simultaneously observe delta bouts in S2. This is relatively important considering that S1 Delta bouts are longer and occupy a larger part of the REM signal. Looking at it the other way around, the overlap was even higher (≈+15.8% overlap, t-test p value = 1e-09). Meaning that the majority of S2 Delta bouts happened at the same time as S1 Delta bouts. In contrast, the distribution of overlaps from either S1 or S2 on M1 Delta bouts is much smaller (respectively -26% and -28%, student pvalue <10-26 for both). Meaning that there was a relatively poor overlap between M1 Delta bouts and those of the somatosensory areas.

Fig.3C shows the different power spectrum profiles for the areas S1, S2 et M1 and during periods of delta in the area S1 during periods of non-delta in the same area. A large difference in S1 was observed between delta and non delta periods (ANOVA p= 2.80e-02, n = 7 mice, posthoc Bonferroni corrected : for S1 p = 2.31e-4) which reflected the efficacy of the method to detect delta periods. However no significant differences in the spectra were found for the area S2 (posthoc, p = 0.056) and the area M1 (posthoc p = 0.1). Since no difference was found for delta we considered the possibility that there is a difference in the power of theta oscillations for these areas. However again there are no significant differences in theta power between delta and non-delta periods for these areas (ANOVA p= 0.5). We then considered the possibility that there is a power difference in delta in areas S1 and M1 for periods of delta and non-delta in area S2. In Figure 3D we see a clear spectral difference for areas S2 and S1 (ANOVA, p = 0.007, n = 7, posthoc for S2: p = 0.003 and for S1: p = 0.038) but not for area M1 (p = 0.07). During these delta and non-delta periods in S2 there was no difference in theta power in the 3 areas (ANOVA, p= 0.5). Finally, we considered the last possibility, i.e. separating delta and non-delta periods in M1. We observed that in this case there is a difference in delta power in areas S1 and S2 (ANOVA p= 0.004, n = 7; posthoc for M1: p= 0.00036; for S1: p= 0.03 and for S2: p = 0.0009). During these delta periods in M1 theta power was not significantly modulated in M1, S1 and S2 (ANOVA, p = 0.059). Thus although the reverse is not true, when there are delta periods in area M1 (which are not frequent, see Figure 2) they tend to occur in delta periods in areas S1 and S2. Overall the analysis of the Delta oscillations suggests that the LFP activity in REM was neither constant nor highly synchronous between the cortical areas.

### 3.2. Computational models of REM sleep in mice

To investigate possible mechanisms to explain these local delta waves during REM sleep, we investigated computational models. We used The Virtual Brain simulation environment (https://www.thevirtualbrain.org/tvb) to simulate the whole brain of the mouse (see Methods). The model was inspired from previous publications (Goldman et al., 2020, 2022) who used mean-field models with conductance-based synaptic interactions to simulate human wakefulness and slow-wave sleep using the TVB platform. Like the human model, the TVB mouse model simulated the asynchronous dynamics reminiscent of Wakefulness (Fig S2 B, Di), and the synchronized slow waves reminiscent of SWS (Fig S2 C, Div). In this model, the transition was driven by modulating the level of adaptation in excitatory populations, which mimics the action of neuromodulators such as acetylcholine, noradrenaline or serotonin, which strongly reduce adaptation in cortical neurons (reviewed in Mccormick (1992)).

In the wakefulness model (W-model), with low adaptation (*b*=5), the neuronal populations are constantly firing with higher firing frequencies for inhibitory cells as observed experimentally (Dehghani et al., 2016). We also see that the different mean-fields are not synchronized together and don’t exhibit clear patterns, thus causing asynchronous dynamics. While in the SWS-model with high adaptation (*b*=60), the neuronal populations alternate between periods of important firing (up to 50Hz) and periods of silence where the mean firing rate fall to 0(Hz), inducing a shift to inverted synchronized slow waves in the LFP with Up and Down state dynamics. We also present the average LFP and spectrum of two intermediate adaptation values for this model. At an intermediate low value (*b* =15nS) most nodes engages in an oscillatory dynamics but it’s less consistent and of smaller amplitude than the oscillations of the SWS-model and there is no real period of silence within the neuronal population. Then, with *b* at 30nS, a value that will be used in our REM model, the simulation is already in a regime similar to our SWS-model but with shorter periods of silence it oscillates faster.

#### Simulating REM sleep

In order to simulate REM sleep activity, we hypothesized that the level of adaptation (or equivalently the action of neuromodulators) was inhomogeneous in the cortex. In Figure 4 we see an example of the heterogeneous distribution of the adaptation parameter b. Motivated by our experimental analysis, we assumed higher values of b in areas such as S1 and S2, and low values in association cortex.

**Figure 4:**
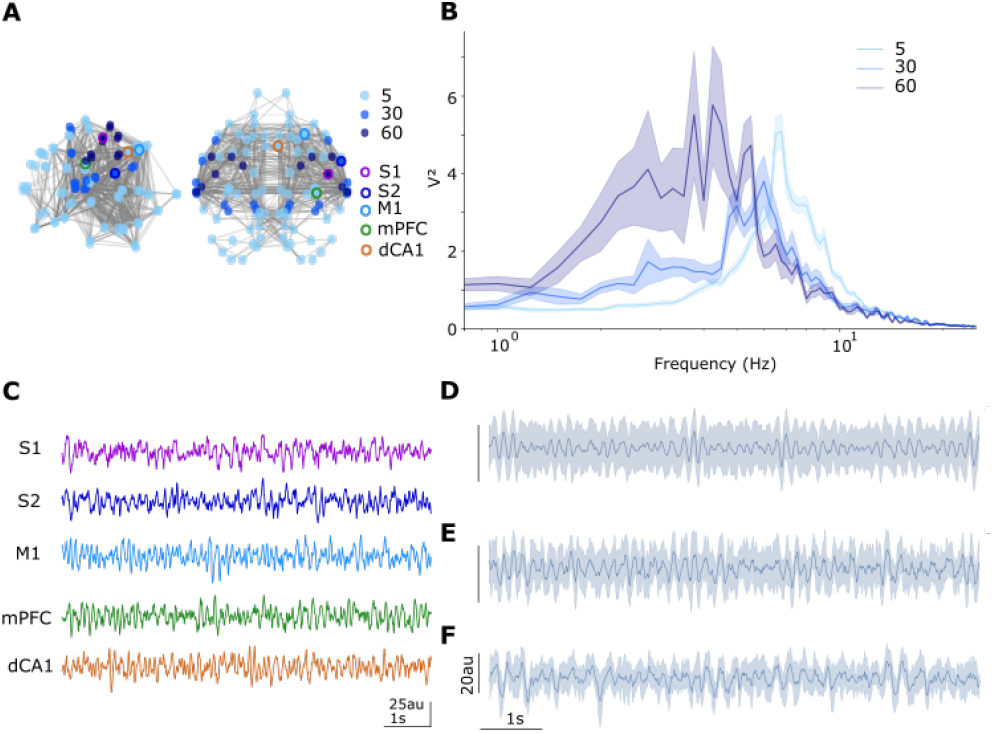
Cortical model with heterogeneous B setting as model of REM. (A) Representation of the Mouse TVB network with nodes for each mean-field. The colour of each node corresponds to the Spike-Adaptation B parameter. Nodes surrounded by a colour circle (Purple, darkblue, lightblue, green and yellow) correspond to the areas studied in figure S7 from the biological dataset. (B) Mean power spectrum for each B parameter value in the model. Power spectrum is computed for each node LFP (with a time window of 4 sec and an overlap of 2 sec), and then averaged across nodes with the same b value. Averaged power spectrums are shown with the SEM. (C) Example 8 seconds of LFP signals from the REM model nodes highlighted in A. (D) Mean LFP signal and its deviation obtained at nodes with *b*=5nS. This corresponds to the adaptation value of the AI reference model. (E) Same as D for nodes with *b*=30nS. (F) Same ad D for nodes with *b*=60nS. It corresponds to the adaptation value of the Up and Down reference model

This type of distribution produced the spatial patterns of local delta waves in the simulated REM sleep. In Figure 4B, the nodes with high adaptation display spectrums dominated by the power between 2 and 6 Hz, which could correspond to the model equivalent of the Delta band. It should be noted that there is an important contrast between this relatively broad power in the Delta band and the strong and sharp peak that the SWS-model produces (Fig S2). For nodes with intermediate level of adaptation, although the PSD peaks at a faster frequency, about 6Hz, there is significantly more power in the low frequencies (below 6Hz) compared to nodes with low adaptation.

In the simulated LFP example of the node corresponding to S1, with high adaptation (Fig 4.C), we can see some slow fluctuations in the signal. This activity is not sustained along the whole simulation but few consecutives cycles could be observed around the 7th second of simulation.Those slower oscillations are of higher amplitude than the portions of the signal that oscillates faster. This slow activity is the model equivalent of the Delta activity we described in the previous analysis. It is harder to find such well defined Delta activity in the simulated LFP for S2. However we can see that the signal of nodes with intermediate level of adaptation, such as the node corresponding to S2, oscillates on average slightly slower and in a slightly more synchronized way than nodes with low adaptation (Fig 4.D,E).

In order to grasp the importance of the topological setup for the spike-triggered adaptation we propose a control model: REM-shuffle, with a random organisation of the parameter *b*. An example is shown in Fig S3 (D,E,F). As expected each REM-shuffle present different global activity and the nodes with high *b* do not always present high slow frequency power (<5Hz). In the example we present, nodes with high *b* peaks at faster frequencies, but some other REM-shuffle (data not shown) could produce dynamics more biased toward slower oscillations than our REM-model. The main point of this control is to show that the adaptation parameter alone is not sufficient to generate the dynamics we want to produce.

Our REM-model, with heterogeneous parametrization of the adaptation present some sporadic slow activity in the nodes with high adaptation that is close to the Delta oscillation observed experimentally. Then we see that areas with intermediate level of adaptation oscillate slower than nodes with low adaptation of the reference model of wakefulness, and could be prone to produce a few delta events.

#### Stimulation in REM model

Finally, to investigate the possibility that Delta in primary sensory cortex impair information integration and thus decreased responsiveness to sensory stimuli (Tononi and Massimini, 2008; Funk et al., 2016), we used our model to compare the responsiveness of different brain states with respect to external inputs (Fig. 5; see also Fig. S5).

**Figure 5:**
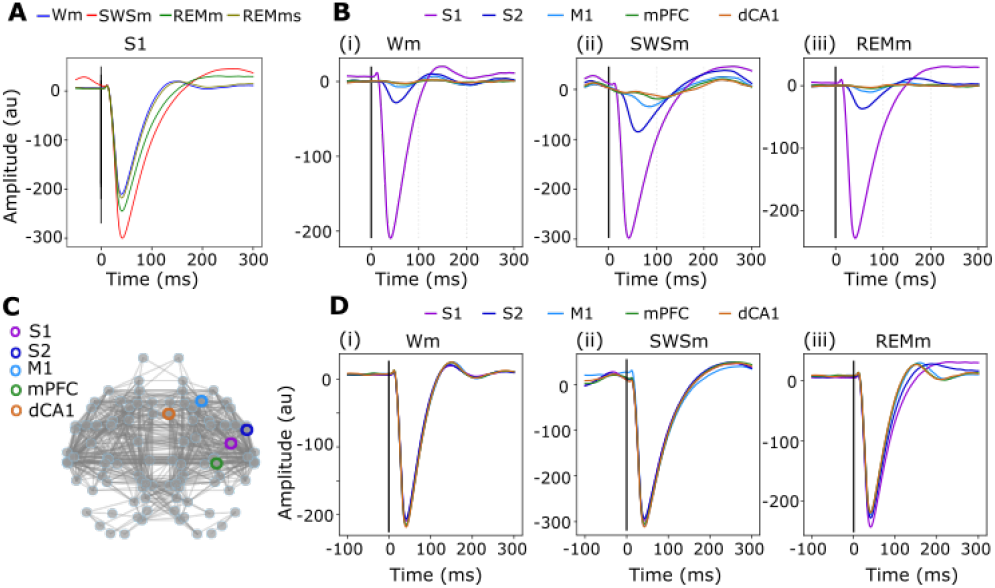
Model response to stimulation in the different states. (A) S1 barrel response when stimulated in the different states (Red corresponds to up and down, Blue to Asynchronous Irregular, Green to REM-model and olive is REM-shuffled). (B) Cortical response to S1 stimulation in the different states. The average LFP response to stimulation in S1 is observed in the different areas of interest across the model states (i: W-model, ii: SWS-model, iii: REM-model). (C) Representation of the model network. Surrounded in colour the five nodes subjected to the stimulation experiment. (D) Response of cortical areas when stimulated across the model states. The colour code corresponds to the areas highlighted in panel C. Each line is the average response observed in an area when it’s stimulated. Each sub-panel corresponding to a different state of the model (i: W-model, ii: SWS-model, iii: REM-model)

Stimulus response can be described as composed of two main parts. The first part is the important and fast negative deflection that occurs in less than 50ms after the stimulation. We can see in Figure S5 (and Fig 5,A), that this initial deflection is of higher amplitude in the SWS-model compared to the W-model. Then in a second time the LFP will return to the baseline level. In the W-model this return is fast and accompanied with a second much smaller negative peak in the LFP. In the SWS-model this second part is slower and is associated with a positive peak around 280m. This reflects a post-stimulus suppression of the neuronal activity. Although the amplitude of the first peak response is exaggerated in our model and the response is a bit slower, the general kinetics are very similar to what could be observed experimentally (Nir et al., 2015; Le Merre et al., 2018). In our REM model, the response to stimulation in areas with high adaptation such as S1 is a bit different to what we saw for the W or SWS models. The amplitude of the first peak is between the two other model levels, although closer to W-model. Then, with a slow hyperpolarized return to the baseline, although without a clear positive peak, the second phase of the response is closer to what we observed in the SWS-model.

In Figure 5.B we can see in our different model states how the response to S1 stimulation propagates across the cortex. We can see that the response in the non-stimulated nodes is much smaller, when there is a clear response. For S2 we can see that the response follows at a smaller scale the dynamics seen in S1 for the W-model and SWS-model. However for the REM-model the second phase of the response in S2 is closer to a slower version of what we see in the W-model, with no clear post-stimulation hyperpolarization. For M1 and the other areas, the response is not clear except in the SWS-model because of amplitude issues. However in the SWS-model the response of M1 peaks later than for S2 and with a smaller amplitude. Such observation could be guessed for the two other states and it follows the response propagation pattern observed in the barrel cortex of awake mice (Le Merre et al., 2018). We do not show stimulation responses in our control REM-shuffle for clarity, but when averaged over multiple simulations (therefore multiple random parametrization) the results are very close to what we observe in the W-model.

Finally we investigated whether our REM-model responds to stimulation in the same way when nodes with different adaptation values are stimulated. As we can see in the Figure 5.D, in homogeneous models such as the Wm or the SWS-model, the response to stimulation in the different areas is extremely similar.On the other hand, in our REM-model we can see that each of our adaptation values elicit a slightly different response. The high adaptation values have the largest first peak and the slowest second phase with marked hyperpolarization. The nodes with intermediate adaptation such as S2 have a faster and more pronounced return to the base line in the secondary part of the response, but it is still relatively slow and there is no negative peak there. And then, the nodes with low adaptation respond to stimulus similarly to the W-model, with the second part of the response as a fast return to the baseline accompanied with a second negative peak.Those results exhibit the variability in stimulation responses one could expect when stimulating during REM, with some areas closely matching the W-model while others have a larger initial peak response and an hyperpolarized second phase as observed in SWS.

## 4. Discussion

In this paper, we have analyzed and modeled cortical REM sleep activity in mice, and in particular the occurrence of slow wave activities. The analysis showed that the REM SWA was mostly constituted of delta oscillations in the 2-4 Hz frequency range. This analysis also shows that REM SWA differed from the SWA of SWS. On a spatial scale, REM sleep expresses Delta waves only in some cortical areas but not in others, in contrast with SWS. On a temporal scale, again in contrast to SWS, REM Delta bouts alternate with periods of asynchronous activity. We next investigated a computational model of the mouse brain build assuming that the neuromodulatory drive, expressed with the adaptation parameter, is heterogeneous in REM sleep. The model further showed that assuming such a heterogeneous neuromodulatory drive, there can be a coexistence between Delta activity and asynchronous activity in the same brain. Below we relate these findings to previous work and delineate possible perspectives to follow-up on this work.

### Experimental analysis of REM sleep slow-waves

The occurrence of local slow-waves during REM was reported in humans and mice (Baird et al., 2018; Bernardi et al., 2019; Funk et al., 2016; Soltani et al., 2019), but our work reveals important differences with the work of Funk et al. First, REM slow-wave activity is significantly faster than that of SWS and is mostly made of Delta activity (2-4Hz), and not SO (<2Hz). It reinforces the idea that SWS SWA is the association of SO and Delta waves (Amzica and Steriade, 1998; Kim et al., 2019) and proposes that, contrary to SWS, REM Delta(2-5Hz) is relatively dissociated from slower activity (<2Hz). Second, it was claimed that REM slow-waves occur only in primary sensory areas and therefore not in secondary or association areas. The clear occurrence of delta oscillations in S2 during REM (Fig. 2) appears to contradict this statement. Moreover, while we found clear evidence of Delta activity in S1 and M1, we did not find it in the primary visual cortex (V1) or and in the primary auditory cortex (Au1, data not shown). Thus, it seems that the results of Funk et al. in visual cortex cannot be generalized to all sensory areas. In these areas theta oscillations were prominent during REM sleep. This topological organisation is in line with the multisite LFP recordings of Soltani et al. (2019). These results suggest that LFP recordings in the neocortex result from a competitive combination of two oscillation generators, one close to the hippocampal theta and the other to the delta prominent in some cortical areas. A recent two-photon experiment by Dong et al. (2022) even propose two sub-stages for REM, a quiet and an active REM sleep, which might correspond to what we characterized as periods with important cortical delta activity in REM, or its absence. Such differences of Delta power in V1 compared to S1 is also observed in human in the REM episodes at the end of the night(Baird et al., 2018). It is also likely that one of the multiple experimental differences (chronic vs acute recording, silicon probes vs tungsten LFP, the detection methods…) could explain those apparently opposite results.

A major structure not yet studied is the thalamus which is known to play a key role in SWS oscillations (Amzica and Steriade, 1998). It is indeed likely that the thalamus, connected to the different sensory modalities, dictate in an essential way the alternation between delta and desynchronized activities. Knowing about thalamic dynamic in REM would help us to understand the origin of this activity, but would be also essential to produce a better, more complete, computational model. Unpublished work by Nadia Urbain’s team (CRNL, Lyon), using intra- and extracellular recordings, shows that the activity of relay neurons in thalamic nuclei connected to S1 and S2 areas changes during REM episodes in a coordinated manner with the LFPs of these cortical areas. Interestingly, in epileptic humans recorded by intracerebral EEG, the thalamic medial pulvinar nucleus is characterized by stable continuous delta waves in REM, yet sometimes interrupted by short periods of rapid activity (Magnin et al., 2004).

One of the essential highlights of Funk et al study is the large neuronal silence observed in the superficial layers of the sensory cortex during REM SWA. This silence period would participate in the decreased responsiveness of REM by making the cortical areas less prone to integrate inputs (Funk et al., 2016), but we saw that REM delta is not constant and alternates with periods of asynchronous activity which may reflect of a cortical state of increased neuronal excitability. The hippocampal activity might be coordinated with the alternation of Delta bouts periods of different activities. Our results suggest that phasic theta often occurs at the beginning and/or end of Delta bouts and might thus participate in the switch from Delta/non-Delta bouts. It has been proposed that phasic theta may be a window of intense synchronization correlatingly activating populations of neurons in the hippocampus and possibly in the neocortex during brief periods of REM. Thus, phasic theta periods during REM sleep could also correspond to short windows of increase in neuronal activity in cortical sensory areas.

The presence of REM SWA in some cortical areas could also be related to a heterogeneous temporo-spatial distribution of neuromodulatory signaling in the neocortex or thalamus. In classical study the unique neuromodulation of REM is associated with elevated Acetylcholine (Ach) (Jasper and Tessier, 1971; hee Lee and Dan, 2012). But it is shown that SWA occurs when cholinergic signaling in S1 is blocked during wakefulness (Eggermann et al., 2014). Therefore if we consider neuromodulation as global, slow and tonic, in REM, the high Ach should prevent SWA. But, with recent technical advances using no longer microdialysis but choline microsensors, Lustig and Sarter(Sarter and Lustig, 2020) suggest in contrast that neuromodulatory signalling is phasic, short duration and very local (synaptic). This conceptual shift help to relate the overall high Ach with the Delta bouts we observed. We hypothesize that in REM the cholinergic signalling toward the somatosensory and motor cortex, or the thalamic nuclei connected to these areas, is low and short duration compared to other brain regions.

A first study is to determine the neocortical-hippocampal relations in REM sleep, and in particular with respect to SWA. One should determine how the hippocampus activity is coordinated with the neocortical activity in REM sleep. Moreover, thalamic activity is clearly an important missing link in determining which structure – hippocampus or thalamus – plays the essential role in the temporo-spatial distribution of cortical activity during REM. During REM the sensory activation threshold is higher, especially for the sense of touch. One may therefore wonder whether SWA in S1 and S2 do not play a role as a sensory filter.

However this should be questioned with respect to the thalamic transmission to sensory cortices in REM. Knowing if the thalamus transmits sensory stimulation similarly as in W is a key to assess how much cortical delta is really about decreasing the responsiveness.

Another follow-up would be to understand the relation between the different transient events that occur during REM, which concern the whole body.REM sleep remains a global phenomenon affecting the whole individual. Multiple organs work together and many factors, such as natural whisking, respiratory or cardiac rhythm could potentially affect brain oscillations (Crochet and Petersen, 2006; Heck et al., 2016; Bergel et al., 2018; Tort et al., 2018b,a).

More experiments could be planned as the cortical activity in REM has received little attention since the transition from cat to rodent as a biological model and is often discarded as just ‘similar to wake’. We also lack specific computational models to question and understand the phenomenon we discussed. With the recent advances, models of W and NREM become more sophisticated and can span multiple scales. New dynamical model of the brain in REM could help to integrate the complex and heterogeneous cortical oscillations we discussed into a larger theoretical framework that could be questioned for new insights. One such model is discussed below.

### Computational models of REM sleep slow waves

The occurrence of local SWA was found before in humans (Baird et al., 2018; Bernardi et al., 2019) and mice (Funk et al., 2016; Soltani et al., 2019). Based on those observations we propose a new model of the whole cortical dynamics of REM sleep. It was built to relate the Delta waves observed in our analysis paper as well as the few previous observations of the phenomenon, and to this regard it is the first of such models. We also provide a randomized control showing that the emergence of REM-sleep features is not due to fine tuning of a series of mean-field models, but is an emergent property of the network. In the model, we could explain the delta waves by assuming a moderate neuromodulatory drive (using an heterogeneous adaptation) in some areas such as S1, S2, which then produce SWA at around 4 Hz. But those Delta waves we see in S1 are of small amplitude. We do not have Delta activity in M1, meaning that with such parametrization, the drive from the somatosensory cortex is not sufficient to produce Delta waves in the LFP. However, surprisingly, the temporal pattern of the Delta activity in S1, with Delta bouts alternating with faster acitivty, seems to be relevant compared to the experimental data although we do not tune the model looking for it. This could be studied more by analysing the delta cycles of an extended dataset of REM model simulations with an appropriate detection.

Our model was used to predict the response to sensory stimulation in the different states. While there are some issues with the responses we observed in the model (addressed later in discussion), the general dynamics of the response across the cortex is consistent with experimental data (Nir et al., 2015; Le Merre et al., 2018).In the REM-model, the responsiveness was modulated with the level of adaptation, evolving from similar to W-model for areas with low adaptation to somehow closer to SWS-model responses for the low to high adaptation levels respectively. In those areas, such as in S1, the peak response was smaller than in the SWS-model which might indicate that a smaller portion of the neuronal population is recruited for the response. But the slow hyperpolarized second phase of the response is very close to what is characteristic of the SWS-model second phase. It comforts the idea that in REM, when we stimulate an area that can present Delta oscillation such as S1, we should observe large peak response followed by an hyperpolarization that disrupts the propagation of activity. This support the idea that REM SWA lead to an impairment of information integration and thus decrease responsiveness to sensory stimuli (Tononi and Massimini, 2008; Funk et al., 2016).

Such a prediction could be tested experimentally by applying whisker stimulation during REM sleep while recording multiple areas. But it should be noted that the stimulus we used in the model is not as natural as a whisker stimulation or a noise presentation done experimentally, but would be closer to TMS experiments such as those performed in humans (Massimini et al., 2007, 2010). In the latest article, authors perfomed TMS on the rostral part of the right premotor cortex during REM sleep. They report a response composed of two peaks close to what they see in wakefulness, but the first peak having a larger amplitude and the second peak a slower amplitude (Massimini et al., 2010). In our data, M1 behaves very similarly in W-model and REM-model. But when compared to the mPFC or dCA1 we could notice a slight tendency toward a slower dynamics in the second phase. We already reported that in our model M1 lack of delta activity compared to the biological data, but it seems likely that with some minor tuning for this node our model would be in accordance with such human results.

The model suffers from a number of limitations. First, it is a population model that cannot explain and detail the activity at the single-neuron level. Therefore such an approach cannot be easily related to experimental data at the cellular level but is more appropriate to relate to population-level variables, such as LFP, EEG or imaging. Second, we pointed to some differences between the model and the experimental analysis. The low Delta activity in S2 and its absence in M1 should be overcomed with a larger area that encompasses M1 for the application of adaptation and some small tuning of the adaptation values used (using more different values seems an interesting approach from some testings, however it makes the model complex to examine). Third, we also pointed to some limits regarding the responses to stimulation. Indeed we have a very large response amplitude and a slightly slower dynamics. While the first issue might be addressed with a refinement of the stimulus used, the second would require to interact with the parameter of the temporal integration of the model, which would in turn require a complete verification of the different parameters used in the model.

The REM model we proposed can be improved in many ways. The topological parametrization of adaptation was kept simple and some issues could be addressed with finer tuning. But one could also think about more complex parametrization of the model that would take into account other aspects of REM sleep such as the characteristic theta activity (Buzsáki et al., 2003; Montgomery et al., 2008) in the hippocampus or the very active prefrontal areas (Renouard et al., 2015; Maciel et al., 2021). Also, the connectome we used in this model remains relatively limited. We used it for multiple practical reasons (easier to test, faster to run), but it lacks multiple brain regions, some of which, like the thalamus, might be critical to understand cortical delta oscillations or stimulus-response. Larger connectomes are now available with more regions and finer parcellation, but correctly integrating new regions according to this newly characterised phenomenon will require a large amount of research and the perspectives of this work are multiple.

First, one could perform a similar approach based on the human brain. One of the problems to overcome is that local slow-waves may not appear in the EEG, and would ideally require intracranial recordings (Magnin et al., 2004). However, such recordings are usually rare in primary sensory areas, so one could use the model to simulate local slow-wave activity, use models to calculate the EEG and determine how such local slow-waves affect the EEG. The aim could be to find identifiable EEG signatures of local slow-waves in REM sleep.

Another possible outcome would be to use intracranial recordings in monkey to confirm the presence of slow-waves in some sensory areas, and test the response to stimuli. A TVB model of the macaque monkey brain is currently available based on the Cocomac database (Shen et al., 2019). One could use this model constrained by intracranial recordings in monkey to further investigate the role of local slow-waves with respect to sensory or other stimuli in REM sleep.

A final, but not least follow-up would be to test the hypothesis behind our REM model: that the occurrence of Delta waves in REM is related to an heterogeneous level of neuromodulation. Because techniques now exist to measure cholinergic release locally (Santos et al., 2015), this question could be addressed experimentally. It would also be possible to use such data to construct a simulation where the level of adaptation (or cholinergic drive) is included not only spatially but also temporally, constrained by such data, leading to more realistic models of state transitions in brain activity during REM sleep.

## Code Availability

The code used for the simulations presented in this paper is available under request and will be available online after publication.

## Acknowledgments

We want to thanks the researchers involved in the ANR PARADOX for their various inputs to the project.

## Supplementary figures

**Figure S1:**
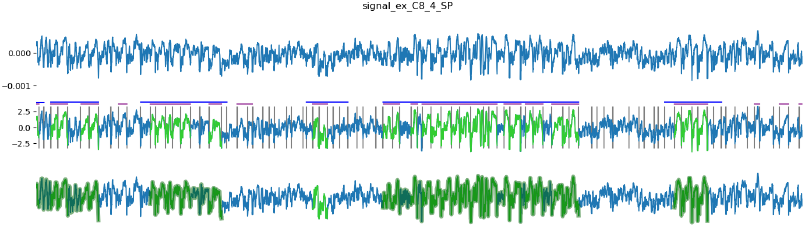
A: raw signal. B: Delta cycle detection, first the signal is segmented into cycles, first threshold detection (blue), second threshold detection (purple). In green are the validated delta cycles C: Delta bouts are shown in darker green)

**Figure S2:**
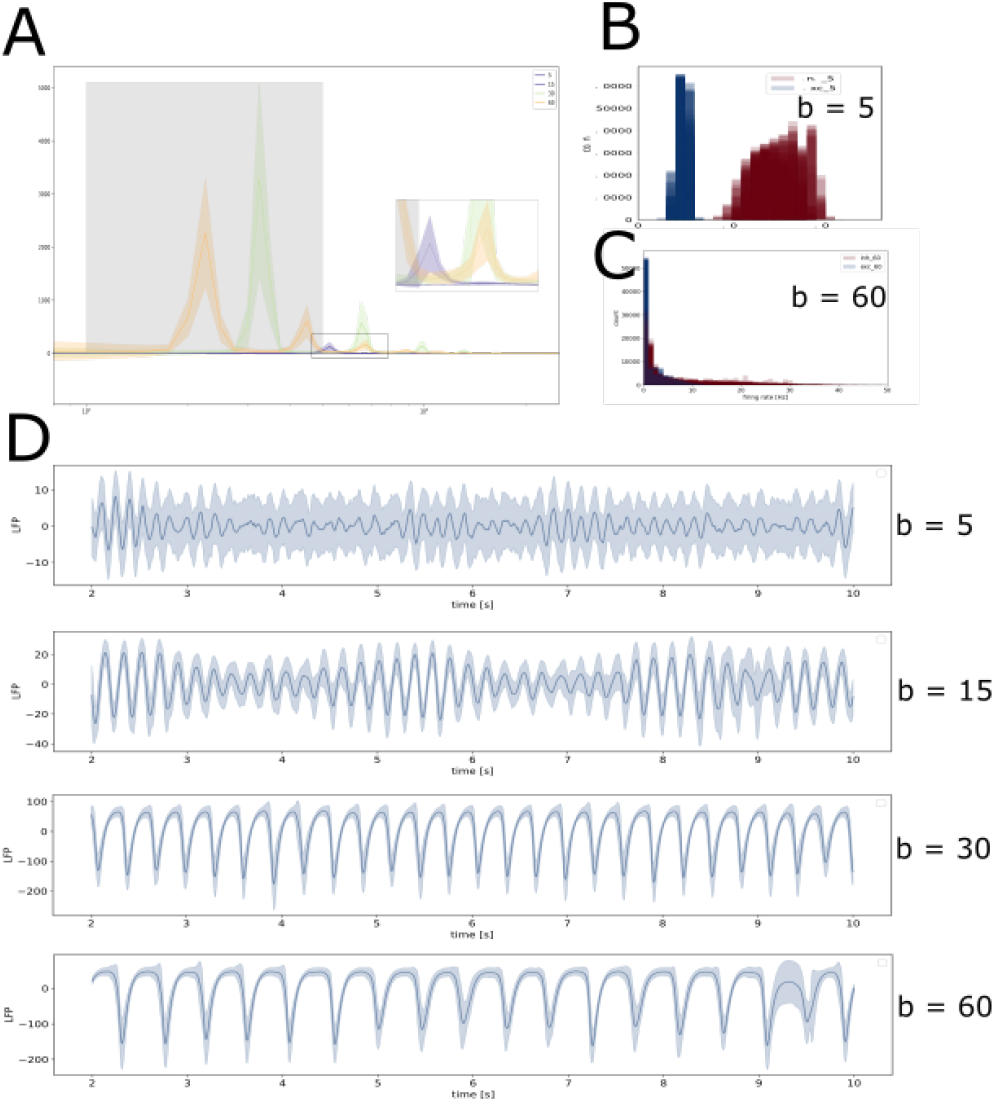
Classical model results with homogeneous adaptation (A): Power spectrum for the four adaptation values presented in D (B): Histogram of the mean firing rate at each nodes for excitatory and inhibitory populations (in Blue and Red respectively) for a W model with b=5. (C): Same as B but for the SWS model with b=60. (D): Mean signal and its deviation for the different value of adaptation discussed.

**Figure S3:**
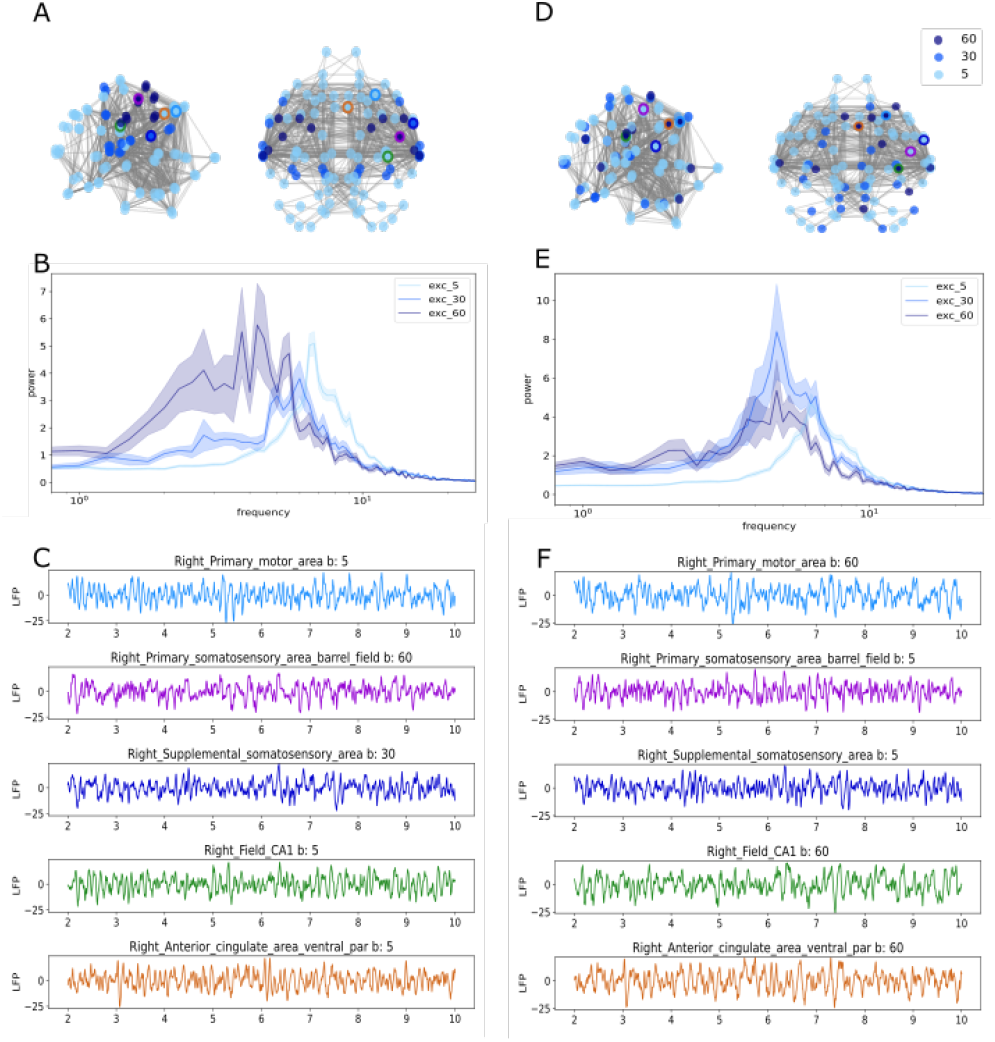
Heterogeneous REM-model vs REM shuffled. Left and Right panels represent the same aspects in two distinct models; the heterogeneous REM-model and its randomised equivalent the REM shuffled. (A/D): Topology of the adaptation parameter in the model. While a clear organisation can be seen in A, it is not the case of D where the setting of b is randomised. (B/E): Power spectrum of the three populations with different adaptation in each model. There is the same number of nodes that participate in the different populations of the two pannels. (C/E): Example LFP signals in the two models. As we can see in F, the b values of the sites of interest are randomised.

**Figure S4:**
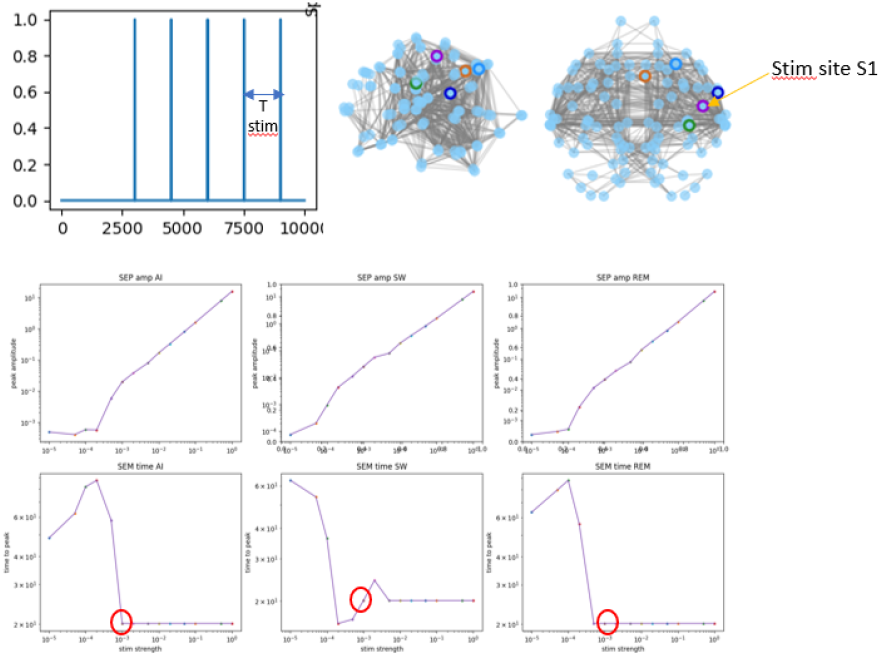
Method of stimulus experiment in the model. (A): Representation of the temporal repartition of the simulations in the protocol. Each stimulus lasts 20ms and is repeated every 1.25±0.25 seconds. (B): Peak amplitude of the Stimulus Evoked Potential (SEP) according to the strength of the stimulus (connection weight). On the Y axis is the amplitude of the response peak observed in the stimulated region. On the X axis is the connection weight between the stimulus source and the stimulated region. Both axis are shown in log. The three panels show the responses values in the different model state (W-model, SWS-model, REM-model from left to right). (C): Time to the peak of SEP according to the strength of the stimulus (connection weight). On the Y axis is the time to the first peak response observed in the stimulated region. On the X axis is the connection weight between the stimulus source and the stimulated region. Both axis are shown in log. The three panels show the responses values in the different model state.

**Figure S5:**
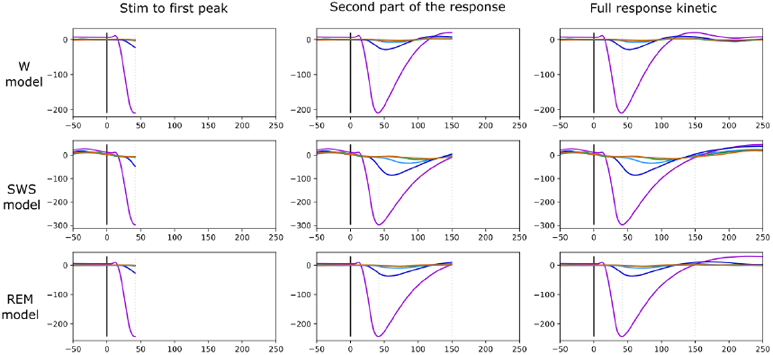
Kinetics of the response to stimulation. This figure shows the temporal evolution of the model in response to stimulation in the three different states. Each line shows one of the states and each panel shows one period manually selected: Panel 1: Response deflection. This panel shows the first part of the stimulus response, up to 42ms after the stimulation, we can see the large deflection of the signal in S1. Similar in duration, the amplitude vary according to the state. Panel 2: Second part of the response. This panel shows how the first response end as well as the the speed to the return to ‘averaged’ value is dependant of the model state. Panle 3: Complete picture of the response observed in the different areas.

**Figure S6:**
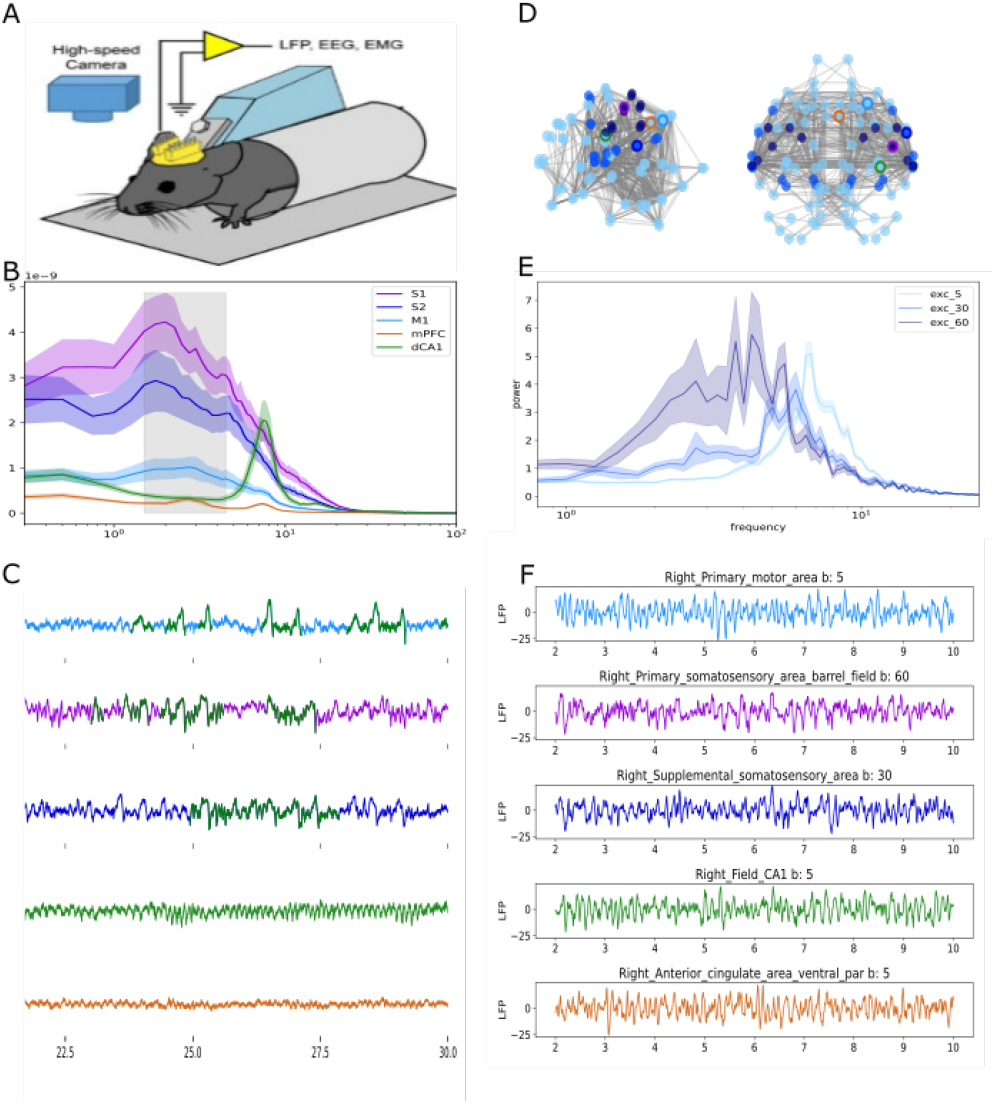
Dynamical response to stimulation in S1. This figure shows the different results of the analysis and the model presented in the article. (A/D): Representation of the data presented in the colomn below. (B/E): Power spectrum observed for the different areas/adaptation values in the corresponding dataset. (C/F) Signal example showing the shape of LFP signal in the biological and simulated data

**Figure S7:**
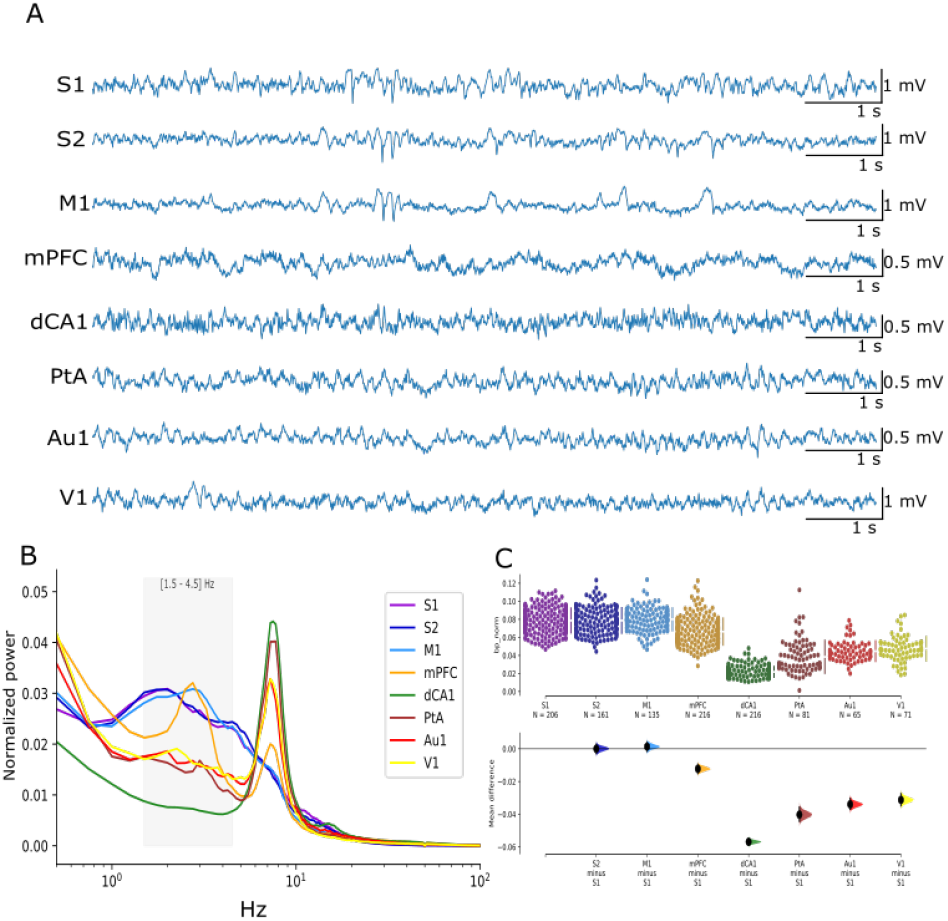
Cortical LFP in REM. This figure shows the activity observed across the cortex during REM. A: 10 sec examples of LFP signals at the different recording sites. B: Normalized power spectrum. C: Comparison of normalized delta bandpower [1.5-4.5Hz] in the different areas during REM

